# Arabidopsis Apoplast TET8 Positively Correlates to Leaf Senescence and *tet3tet8* Double Mutants are Delayed in Leaf Senescence

**DOI:** 10.1101/2024.05.10.593620

**Authors:** Jayde A. Zimmerman, Benjamin Verboonen, Andrew P. Harrison Hanson, Judy A. Brusslan

## Abstract

Extracellular vesicles (EVs) are membrane-bound exosomes secreted into the apoplast. Two distinct populations of EVs have been described in Arabidopsis: PEN1-associated and TET8-associated. We previously noted early leaf senescence in the *pen1* single and *pen1pen3* double mutant. Both PEN1 and PEN3 are abundant in EV proteomes suggesting EVs might regulate leaf senescence in soil-grown plants. We observed that TET8 is more abundant in the apoplast of early senescing *pen1* and *pen1pen3* mutant rosettes and in older WT rosettes. The increase in apoplast TET8 in the *pen1* mutant did not correspond to increased *TET8* mRNA levels. In addition, apoplast TET8 was more abundant in the early leaf senescence *myb59* mutant, meaning the increase in apoplast TET8 protein during leaf senescence is not dependent on *pen1* or *pen3*. Genetic analysis showed a significant delay in leaf senescence in *tet3tet8* double mutants after six weeks of growth suggesting that these two tetraspanin paralogs operate additively and are positive regulators of leaf senescence. This is opposite of the effect of *pen1* and *pen1pen3* mutants that show early senescence and suggest PEN1 to be a negative regulator of leaf senescence. Our work provides initial support that PEN1-associated EVs and TET8-associated EVs may have opposite effects on soil-grown plants undergoing age-related leaf senescence.

## Introduction

Extracellular Vesicles (EVs) are secreted vesicles surrounded by one or more lipid bilayers produced by bacteria, archaea, fungi, animals and plants (Bose et al., 2020; Dai et al., 2020; Liebana-Jordan et al., 2021; Chaya et al., 2024). Heterogenous EVs are broadly divided into three categories based on size and biogenesis. Exosomes range from 50-150 nm and form as intraluminal vesicles in multivesicular bodies (MVB), ectosomes (50-1000 nm) pinch off from the plasma membrane, and apoptotic bodies (50-5000 nm) form during plasma membrane blebbing. Exosomes play many roles in intercellular signaling within the mammalian tumor microenvironment and in young and aging cells (Lananna and Imai, 2021).

Plant EVs were first noted in barley epidermal cells inoculated with biotrophic fungi. Outward budding paramural bodies were observed by transmission electron microscopy. The paramural bodies localized to the site of cell wall appositions that slowed fungal penetration and stained for H_2_O_2_ and the PRX7 vacuolar peroxidase, suggesting a role in reactive oxygen species (ROS)-related cell wall defense. (An et al., 2006b; An et al., 2006a)

Plant EVs were later purified from apoplast fluid by differential centrifugation (Regente et al., 2009; Rutter and Innes, 2017; Cai et al., 2018). The two most studied plant EVs are described as PEN1-associated and TET8-associated. Both PEN1 and TET8 are plasma membrane proteins. PEN1, also known as SYN121, is a syntaxin while TET8 is a member of the tetraspanin family, a protein component shared with animal exosomes. Tetraspanins CD63, CD81 and CD9, are abundant in purified mammalian exosomes (Jeppesen et al., 2019). PEN1-associated EVs are enriched after a 40,000 x g centrifugation and estimated to be 150 nm in diameter (Rutter and Innes, 2017). TET8-associated EVs show enrichment after 100,000 x g centrifugation (He et al., 2021)). Arabidopsis expressing mCherry-PEN1 and TET8-GFP show two non-overlapping EV populations, but differential centrifugation does not entirely separate these two classes of EVs (He et al., 2021). In addition, the two EV populations may have distinct biogenesis pathways as revealed by partial overlap of TET8-YFP with the MVB marker ARA6-CFP, but no overlap between ARA6-YFP and CFP-PEN1 (Cai et al., 2018; He et al., 2021). However, the PEN1-enriched EV proteome does share components with the ARA6 subcellular proteome (Heard et al., 2015).

The first EV proteome was determined from iodixanol density-gradient purified vesicles (Rutter et al., 2017). Proteins related to defense, ROS, membrane trafficking, vesicle transport and ion transport were identified. A second EV proteome also harbored defense and stress response proteins (He et al., 2021). PEN1 and TET8 were detected in the two proteomes, and in both cases PEN1 was more abundant. Both proteomes had high peptide counts for PEN3, an ABC transporter protein with specificity for defense-related indole-metabolites such as camalexin (Lu et al., 2015; He et al., 2019). The sorghum EV proteome shares many Arabidopsis EV proteins suggesting EV conservation between monocots and dicots (Chaya et al., 2024).

mRNAs reside inside EVs, and two of these, *SAG21* and *APS1*, are transferred to biotrophic fungal cells, where they are translated and contribute to reduced fungal infectivity (Wang et al., 2024). Other EV-resident RNAs are tiny RNAs (10-17 nucleotides) from coding sequences, transposable elements and intergenic regions. The tiny RNAs are mostly derived from the middle of cellular RNAs and likely represent degradation products that remain after 5’ and 3’ exonuclease digestion (Baldrich et al., 2019). Trypsin and RNase A digestion were performed to remove protein-bound RNA co-purifying with EVs. After this treatment, only seven miRNAs were shown to be EV resident with six of these being passenger strands. The function of the EV-resident tiny RNAs and miRNAs has not been determined (Zand Karimi et al., 2022).

Additional genetic analysis suggests EVs play a role in defense. PEN1 enhances defense against fungal haustorium penetration (Collins et al., 2003) and is thought to facilitate vesicle formation for cell wall reinforcement. *tet8tet9* double mutants are more susceptible to *B. cinerea* infection, displaying larger lesions than WT (Cai et al., 2018). In contrast, *tet8* mutants are less responsive to the defense hormone salicylic acid (SA) showing reduced ROS production and cell death (Liu et al., 2020). *tet8* mutants retain ∼40% of EVs; this partial reduction in EVs could be explained by PEN1-associated EVs or EVs with other tetraspanins. *TET9* is the closest paralog of *TET8*, but it is expressed at low levels under the standard growth conditions used in the Liu et al, 2020 study. *TET3* is in a sister clade to *TET8* and is highly expressed (Supplemental Figure 1). *TET3* expression is ABA, drought and cold-responsive (Wang et al., 2015) and TET3 may be able to form EVs in *tet8* mutants.

We have previously reported that the *pen1pen3* double mutant displays early SA-dependent leaf senescence (Crane et al., 2019). Leaf senescence is the gradual dismantling of older leaves that maximizes nutrient export prior to leaf death. Both PEN1 and PEN3 are abundant EV proteins suggesting that EVs may play a role in leaf senescence. In addition, SA has been shown to increase PEN1-associated EVs (Rutter and Innes, 2017; He et al., 2021) and age-induced leaf senescence is regulated by SA (Buchanan-Wollaston et al., 2005). To provide evidence supporting a role for EVs in leaf senescence, we have quantified TET8 in apoplast fluid as an approximation of TET8-associated EVs. We note that TET8 signal is increased in the early senescent *pen1* and *pen1pen3* and in older WT rosettes. Apoplast TET8 is dramatically increased in older rosettes of *pen1* mutants, but not in younger rosettes, suggesting apoplast TET8 is not compensating for loss of PEN1, but is associated with leaf senescence. Additionally, apoplast TET8 is increased in the early-senescent *myb59* mutant (He et al., 2023). Beyond correlation, *tet3tet8* double mutants show a significant delay in leaf senescence and the early leaf senescence displayed by *pen1* mutants is reversed in *pen1tet3* and *pen1tet8*. Our apoplast TET8 and genetic data point to opposing roles for PEN1 and two tetraspanins (TET3 and TET8) in leaf senescence.

## Materials and Methods

### Plant Growth Conditions

*Arabidopsis thaliana* seeds were sowed in Sunshine Mix #4 soil, Sun Gro Horticulture, Agawam, MA and cold stratified at 4^°^C for 5 days before being grown under long-day growing conditions (20 hours of light at 40 µmole photons m^-2^ sec^-1^, 24^°^C). Rosettes were supplemented every week with 10 mL of GRO POWER 4-8-2 (Gro-Power, Chino, CA) per gallon of water and received an application of the larvicide, Gnatrol ® (0.3g per 400 ml), on the soil surface to prevent fungus gnats at the time of seed sowing.

### Mutants

Mutant lines used in this study are listed in Supplemental Table 1. Seeds were obtained from the Arabidopsis Biological Resource Center (ABRC, Ohio State University). Prior to experimentation, mutations or insertions were verified by using primers designed by the iSect tool at SALK T-DNA express (http://signal.salk.edu/tdnaprimers.2.html). Flanking sequences were amplified using a T-DNA border primer and sequenced to identify the insertion site. Point mutants were identified through sequencing after amplification of the desired region. Primers are listed in Supplemental Table 1.

### DNA Isolation

DNA was isolated using Plant DNAzol™ (ThermFisher, Inc.) according to manufacturer’s instructions. *Taq* polymerase (New England Biolabs, Inc.) was used for standard PCR reactions in a Bio-Rad T100 thermal cycler using a 57°C annealing temperature.

### Apoplast Extraction

The protocol was adapted and modified from (Rutter and Innes, 2017) and (He et al., 2021). Rosettes were harvested by cutting plants at the hypocotyl at the soil line and removing inflorescences. Between 5-10 grams (approximately between 12-24 plants) were collected, washed in distilled water to remove soil and then submerged in 200 mL vesicle isolation buffer (VIB, 20mM MES hydrate, 2mM CaCl_2_, 0.1M NaCl pH 6.0 with HCl) inside a 250 mL beaker. The submerged rosettes were vacuum infiltrated at -30 kPa for 30 seconds. The infiltrated rosettes were blotted dry and placed in 50 mL conical tubes punctured at the base. A slow-speed centrifugation (900 x g, 15 min, 4°C) was used to collect the raw apoplast (approximately 3 ml). The raw apoplast was centrifuged (2,000 x g, 30 min, 4°C) to pellet large cellular debris. The supernatant was transferred to autoclaved 1.5 ml microcentrifuge tubes and spun to pellet larger non-EV vesicles (10,000 x g, 30 min, 4°C). The resulting supernatant was considered partially pure apoplast that was used for immunoblotting or stored at -20°C.

### Immunoblotting

The sample preparation for whole leaf extract and SDS-PAGE protocol were adapted from Martinez-Garcia et al. (1999) with 9 µL of whole leaf extract and 1 µL of buffer Z per lane. 14 µL of apoplast and 4 µL of 4X loading buffer (250 mM Tris-HCl, pH 6.8, 10% SDS, 30% glycerol, 10 mM DTT, 0.05% bromophenol blue) were denatured at 95°C prior to electrophoresis. After electrophoresis, protein was transferred to nitrocellulose using a Bio-Rad Transblot Turbo Transfer system. Immunoblotting was performed using the TET8 antibody (PhytoAB PHY1490S, lot# 1941A5) and the goat anti-rabbit HRP secondary antibody (PhytoAB PHY600, lot #2206A5). HRP was detected with Super Signal™ West Pico PLUS (ThermoFisher, Inc.). Relative protein present in each sample was quantified with Coomassie brilliant blue dye using the most prominent band: RbcL. TET8 signal was normalized based on relative protein amount. Imaging for HRP detection and Coomassie staining was performed with a ProteinSimple FluorChem Fluorescent Western Blot Imaging System.

### Real-time qPCR

RNA was isolated from leaves 4 and 5 using Trizol and cDNA was synthesized using MuMLV reverse transcriptase (New England Biolabs, Inc.) primed with random hexamers. *ACT2, NIT2,* and *TET* primers (Supplemental Table 1) at 70 nM with 1X SYBR Green (qPCRBIO SyGreen, Blue Mix HI-ROX, PCR Biosystems) were used to amplify a 1:16 dilution of cDNA in triplicate in a Quantstudio™6 PRO real-time thermal cycler (ThermoFisher, Inc.) with a 61°C annealing temperature. *NIT2* or *TET* gene expression normalized to *ACT2* was calculated (40-ΔCt, (Livak and Schmittgen, 2001; Garapati et al., 2015).

### Chlorophyll

Leaf 3 was collected, weighed, transferred to a 1.5 ml tube, frozen in liquid nitrogen, and stored at - 80°C. Leaves were submerged in 800 µL dimethylformamide (DMF) upon thawing and chlorophyll was eluted into the DMF in the dark at room temperature for 16-24 hours. Absorbance was measured at 647 nm and 664 nm, and chlorophyll concentrations were calculated using equations described in (Porra et al., 1989) and normalized to leaf weight.

Dark-Induced Leaf Senescence Leaves 3 and 4 were collected from plants after 21 days of growth. Leaves 3 and 4 emerge from the meristem close to the same time and are nearly equal in age and size at 21 days. Leaves were flash frozen (day 0) or arranged on filter paper (Whatman Hardened Ashless 125mm) saturated with 1.5 mL of 3 mM MES (pH 5.8) and adhered to a petri dish. Petri dishes were sealed with parafilm and then placed in dark canisters in the growth chamber. Dark canisters were removed after three days, and leaves were flash frozen in 1.5 ml tubes and stored at -80°C prior to chlorophyll measurements. Samples were not normalized to weight as the size of the leaves was uniform.

### Statistical Analysis

All statistical analyses were performed in GraphPad Prism. Outliers were removed using default parameters and log normality was verified prior to analyses. In some instances, data were log transformed prior to statistical analysis. Different experimental conditions were compared to the control using a one-way ANOVA with Dunnett’s correction for multiple comparisons. P-values were reported as < 0.05 (*), <0.01 (**), < 0.001 (***) and < 0.0001 (****).

## Results

### Apoplast TET8 increases in early senescing p*en1* and *pen1pen3*

We have previously reported accelerated SA-dependent leaf senescence in *pen1pen3* mutants. PEN1 and PEN3 are abundant EV proteins suggesting EVs may play a role in the regulation of leaf senescence. To elucidate the relationship between leaf senescence and EVs we measured the amount of apoplast TET8 and quantified leaf senescence in WT, *pen1*, *pen3, pen1pen3* and *tet8* mutants after seven weeks of growth (Collins, 2003, Stein, 2006, Boavida 2013). *tet8* served as a negative control for the TET8 antibody. A significantly brighter apoplast TET8 signal was observed in the *pen1* and *pen1pen3* mutants compared to the wild-type and the *pen3* mutant (Figure 1A-B, Supplemental Figure 2). TET8 was enriched in apoplast fluid, only being detectable when substantial amounts of whole leaf extract were (Collins et al., 2003; Stein et al., 2006; Boavida et al., 2013) loaded on the gel as shown by Coomassie staining. The elevated apoplast TET8 in *pen1* and *pen1pen3* was associated with accelerated leaf senescence. This was indicated by the significant increase in expression of the leaf senescence marker transcript *NIT2* (Figure 1C; Brusslan et al., 2012; Brusslan et al., 2015) and accompanied by significantly reduced chlorophyll (Figure 1D). Taken together our results suggest that *pen1* and *pen1pen3* mutants have elevated levels of apoplast TET8 correlating to accelerated leaf senescence.

**Figure 1.**
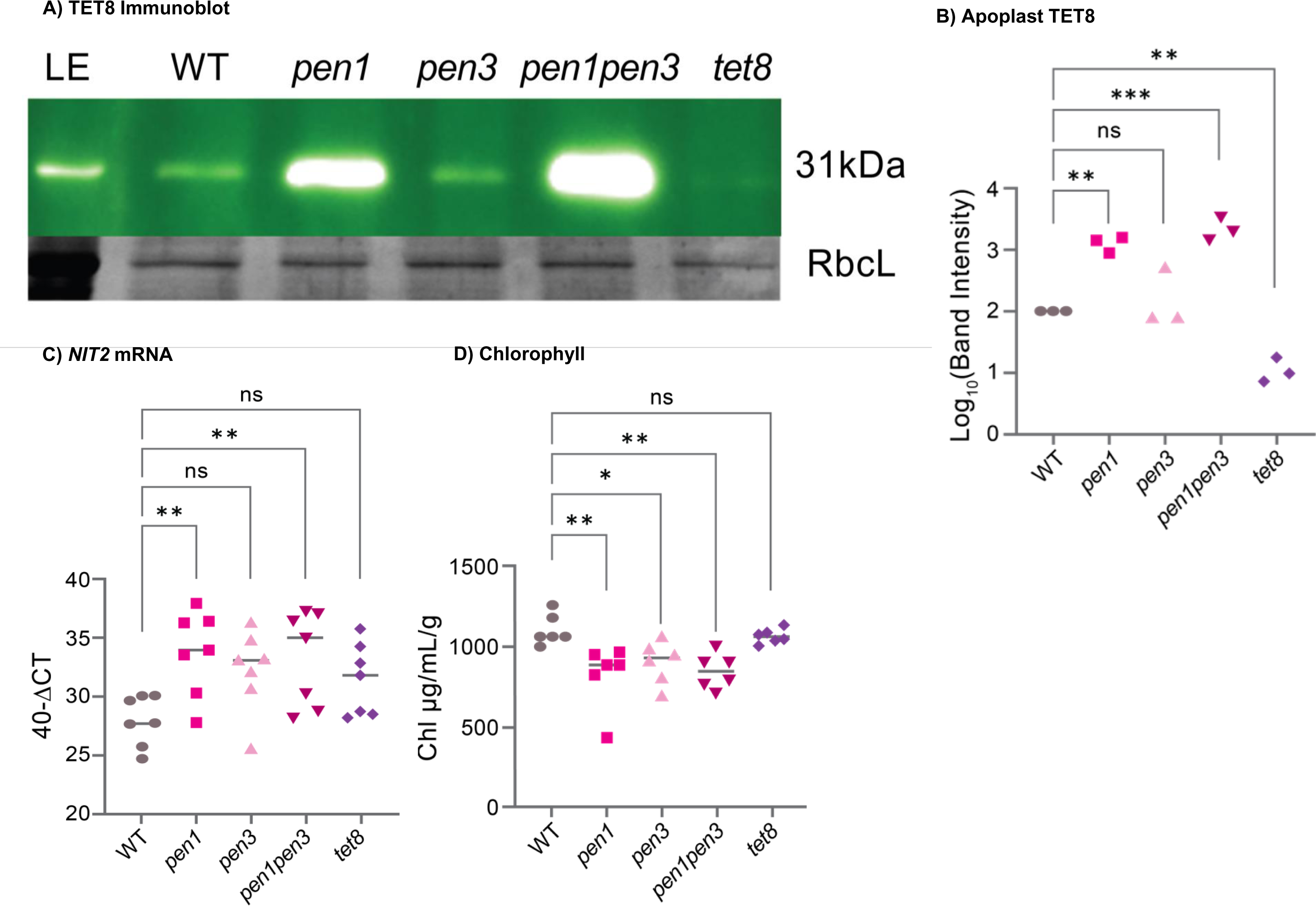
Apoplast TET8 increases in early leaf senescence mutants *pen1* and *pen1pen3*. A) TET8 was detected in leaf extract (LE) from WT and apoplast fluid from WT, *pen1*, *pen3*, *pen1pen3* and *tet8* using an immunoblot normalized to Coomassie-stained RbcL. B) Normalized TET8 chemiluminescent signals from three independent experimental replicates are shown. C) From the same 8-week-old tissue, leaves 4 and 5 were harvested for RNA extraction and *NIT2* gene expression, normalized to *ACT2*, was quantified. D) Leaf 3 was harvested, and total chlorophyll was measured and normalized to fresh weight. A stronger TET8 signal in the apoplast is accompanied by higher *NIT2* expression and lower chlorophyll levels showing a positive correlation between apoplast TET8 and leaf senescence. *tet8* (SALK_136039) served as a negative control for the antibody. Additional experimental replicates are shown in Supplemental Figure 2.

### Apoplast TET8 increases with plant age

To support the correlation between early leaf senescence and apoplast TET8, we next determined if the progression of leaf senescence in WT was associated with an increase in apoplast TET8. We collected apoplast fluid from WT rosettes at different ages and isolated RNA and chlorophyll from the same plants (Figure 2, Supplemental Figure 3). Older rosettes (eight weeks) presented a significant increase in apoplast TET8 compared to five-week-old plants (Figure 2A and 2B). The rosettes were undergoing leaf senescence shown by increased *NIT2* expression (Figure 2C) and a reduction in chlorophyll (Figure 2D). Older rosette leaves were used for chlorophyll (leaf 3) and *NIT2* gene expression (leaves 4 & 5) while whole rosettes were used for apoplast extraction, which may explain why significant changes in leaf senescence markers (6 weeks) occurred earlier than increased apoplast TET8 (8 weeks). These results show that apoplast TET8 increases during the progression of age-related leaf senescence.

**Figure 2.**
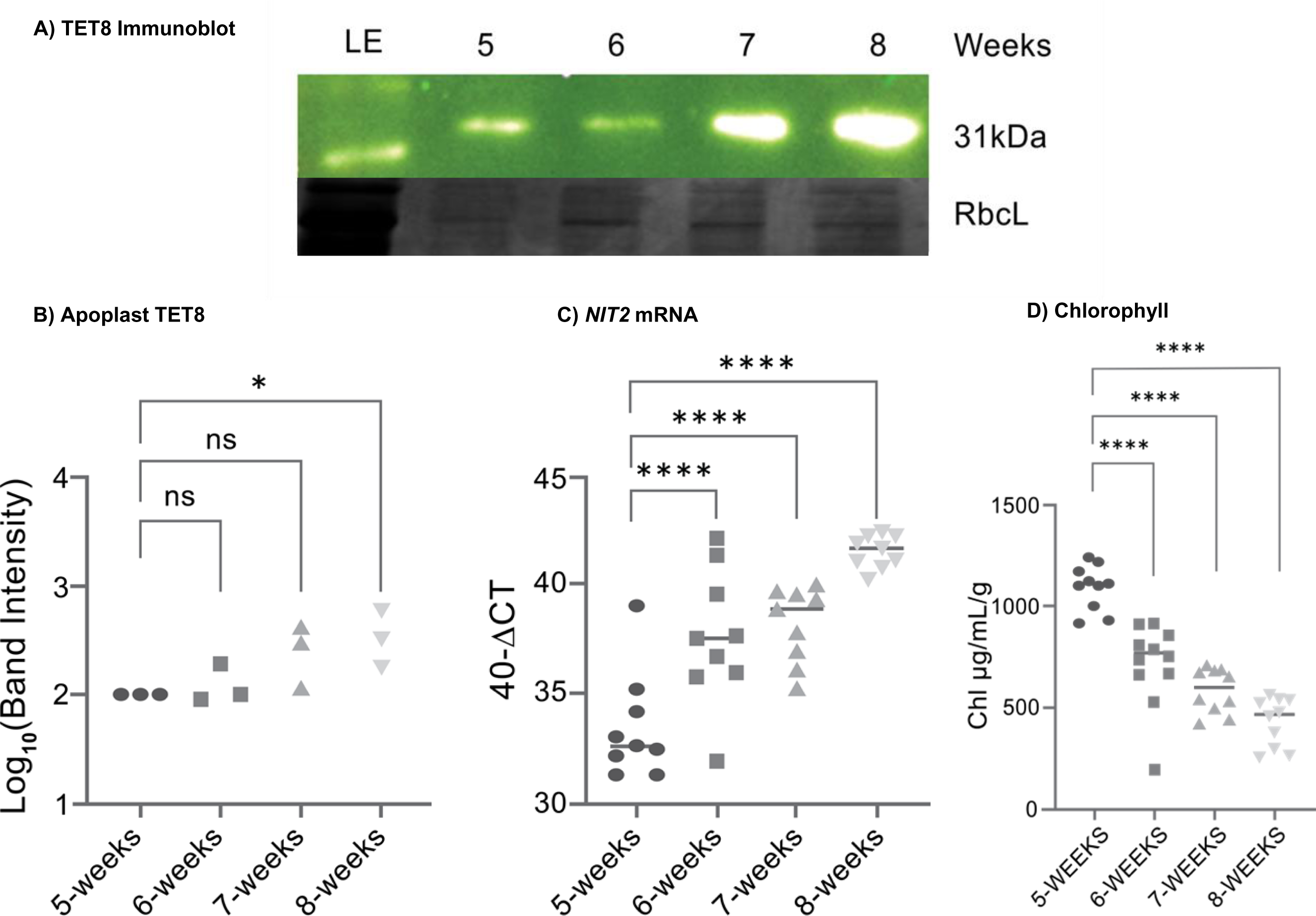
An increase in apoplast TET8 was observed in older WT rosettes. A) Apoplast TET8 was detected from rosettes harvested at 5 to 8 weeks using an immunoblot normalized to Coomassie-stained RbcL. B) Normalized TET8 chemiluminescent signals from three independent experimental replicates are shown. C) *NIT2* gene expression (leaves 4 &5) and D) chlorophyll (leaf 3) are shown from plants grown alongside those used for apoplast extraction. Additional experimental replicates are shown in Supplemental Figure 3.

### Apoplast TET8 in increased in *pen1* mutants only during leaf senescence

Leaves contain both PEN1-associated and TET8-associated EVs, and loss of PEN1 in the *pen1* mutant could be compensated by an increase in TET8-associated EVs, independent of leaf senescence. To explore this possibility, apoplast was isolated from ∼4 g of four-week (non-senescent) and eight-week-old (senescent) WT and *pen1* rosettes. Apoplast TET8 is equally abundant in WT and *pen1* at 4 weeks, but significantly more abundant in *pen1* at 8 weeks (Figure 3A-C, each sample is an independent experimental replicate). The increase in apoplast TET8 is not accompanied by an increase in *TET8* gene expression (Figure 3D) demonstrating that *TET8* mRNA levels do not directly reflect apoplast TET8 abundance. These findings show that apoplast TET8 does increase in *pen1* during leaf senescence, but not before, and that this change does not occur at the level of gene expression. As *pen1* displays earlier leaf senescence, these findings also support the positive relationship between apoplast TET8 and leaf senescence.

**Figure 3:**
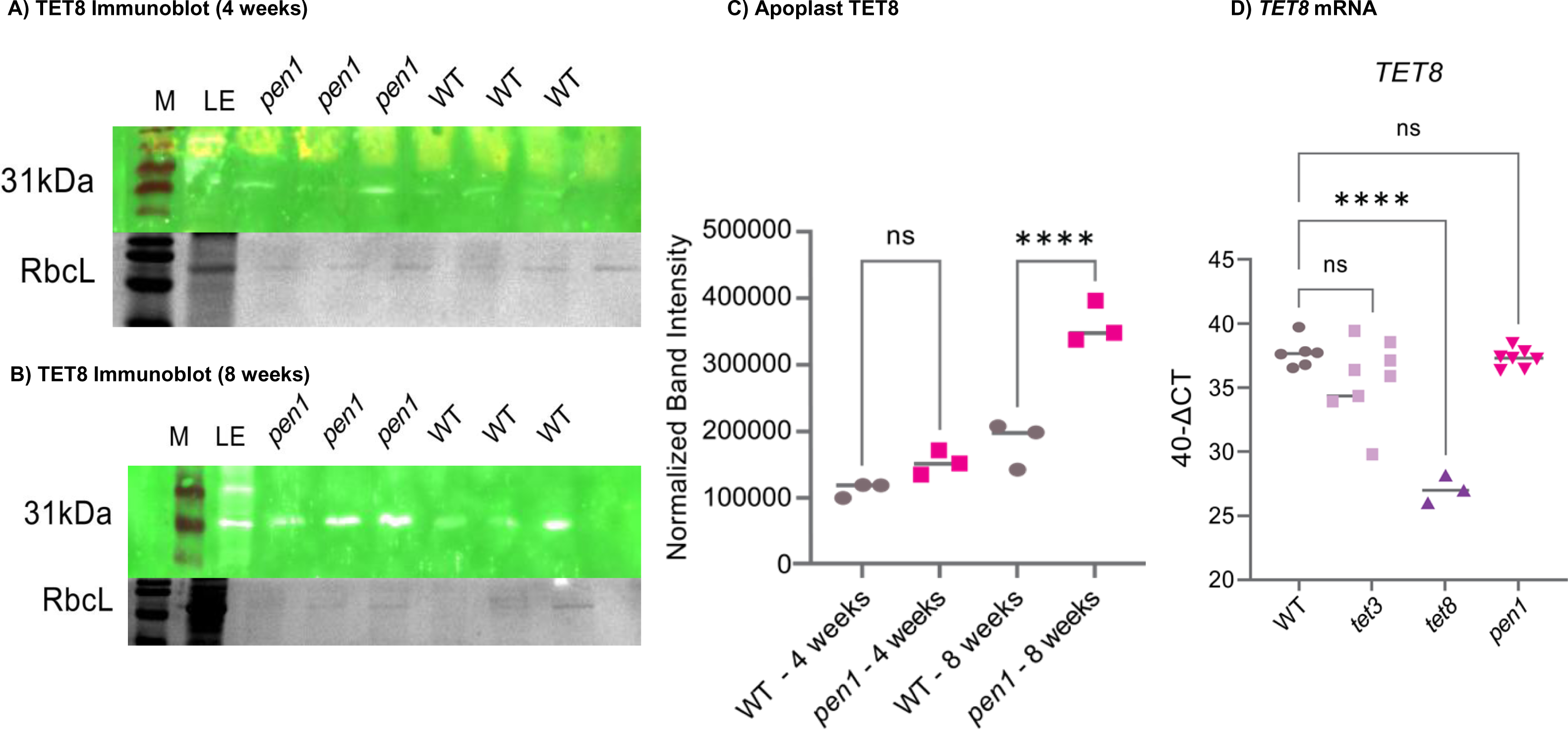
Apoplast TET8 is more abundant in *pen1* mutants during leaf senescence but not in younger rosettes. A) Apoplast was extracted from WT and *pen1* rosettes after 4 weeks **(A)** or 8 weeks (**B**) and subject to immunoblot analysis and normalization with Coomassie-stained RbcL. Three experimental replicates at each age are shown in the immunoblot. **C**) Normalized TET8 chemiluminescent signal from the three replicates are shown and the significant difference after 8 weeks of growth between WT and *pen1* is evident. **D**) *TET8* gene expression, normalized to *ACT2*, was measured by RT-qPCR in leaves 4 and 5 from 8-week-old plants. *TET8* mRNA levels were unchanged in WT, *tet3* (SAIL_617_C05) and *pen1*, but partial transcripts were reduced 1000-fold (2^-10) in *tet8*.

### Apoplast TET8 is increased in the early senescing *myb59* mutant

We were concerned that early senescence in *pen1* and *pen1pen3* might complicate our experiments since PEN1 and PEN3 are EV-resident proteins. Hence, we measured apoplast TET8 in an early leaf senescence mutant that was not in a gene encoding an EV component. We obtained the *myb59* allele (GABI-KAT_627C09, He et al., 2023) and observed accelerated leaf senescence when plants were transferred to higher light intensity for the last three weeks of the six-week growth period (Figure 4A). Early leaf senescence was not observed when *myb59* remained in lower light intensity. Apoplast fluid was similarly extracted from WT and *myb59* (both grown together under the same light conditions), and Figure 4B-C show apoplast TET8 was increased in the early senescing *myb59* in three separate experiments. This supports a positive correlation between leaf senescence and apoplast TET8 accumulation that is not related to perturbation of *pen1*/*pen3*.

**Figure 4:**
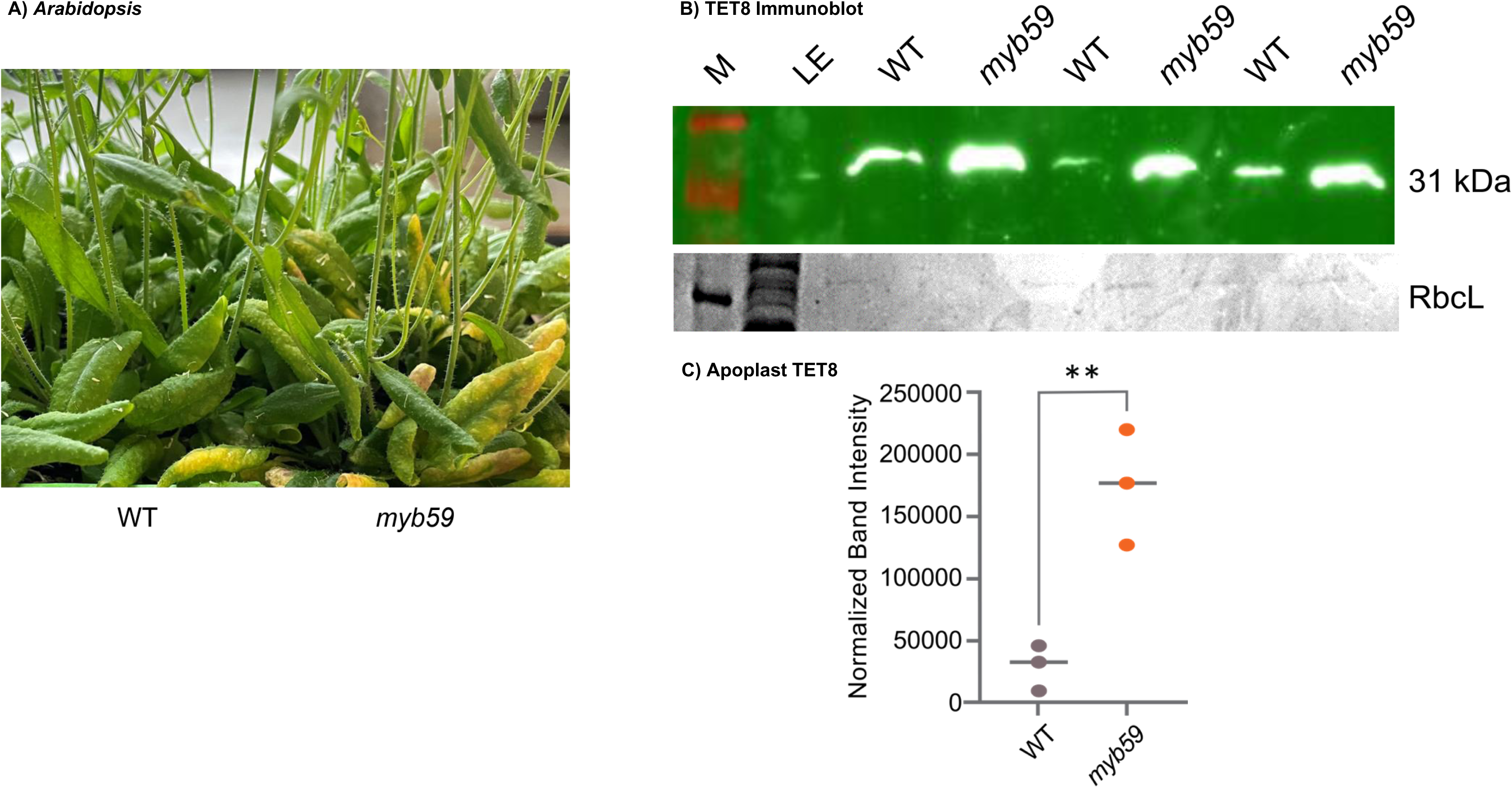
The *myb59* mutant displays early leaf senescence and has higher apoplast TET8. A) WT and *myb59* were grown for 3 weeks at 40 μmol photons/m^2^/sec and then transferred to 160 μmol photons/m^2^/sec for three additional weeks. Reduced chlorophyll in the *myb59* line (GABI-KAT 627C09: T-DNA insertion in exon 3 of AT5G59780) is visible when compared to WT. B) WT and *myb59* apoplast from rosette leaves was subject to TET8 immunoblot analysis. C) Normalized TET8 chemiluminescent signal from the three independent replicates is shown and the significant difference between WT and *myb59* is evident.

### Genetic analysis indicates opposing roles for PEN1 and tetraspanins TET3 and TET8 in leaf senescence

Since EVs are identified as PEN1-associated and TET8-associated, we generated a *pen1tet8* mutant to reduce EV production. We noted that *TET3* is reported to be induced by leaf senescence and its expression level is similar to *TET8* (Supplemental Figure 1, Figure 3D and Supplemental Figure 4B). Although TET3 is not observed in any EV proteome, its abundance and similarity to TET8 suggest it may have related functions. For this reason, we also produced *pen1tet3* and *tet3tet8* double mutants. The *tet3* T-DNA mutant allele was shown to be a strong knockdown (Supplemental Figure 4B). Plants were grown for seven weeks and chlorophyll and *NIT2* gene expression were used to quantify leaf senescence (Figure 5). Unexpectedly, we noted that the early leaf senescence observed in *pen1* (Figure 1C-D) was reversed by *tet3* or *tet8* mutations in double mutants. In addition, the *tet3tet8* double mutant showed a marked delay in leaf senescence. These findings suggest that PEN1 and the two tetraspanins play opposite roles in leaf senescence with TET3 and TET8 promoting leaf senescence (the *tet3tet8* double mutant delays leaf senescence) and PEN1 slowing leaf senescence (the *pen1* mutant accelerates leaf senescence). The double mutants behaved similarly to WT during dark-induced leaf senescence (Figure 6) showing these opposing roles are specific to age-related leaf senescence.

**Figure 5:**
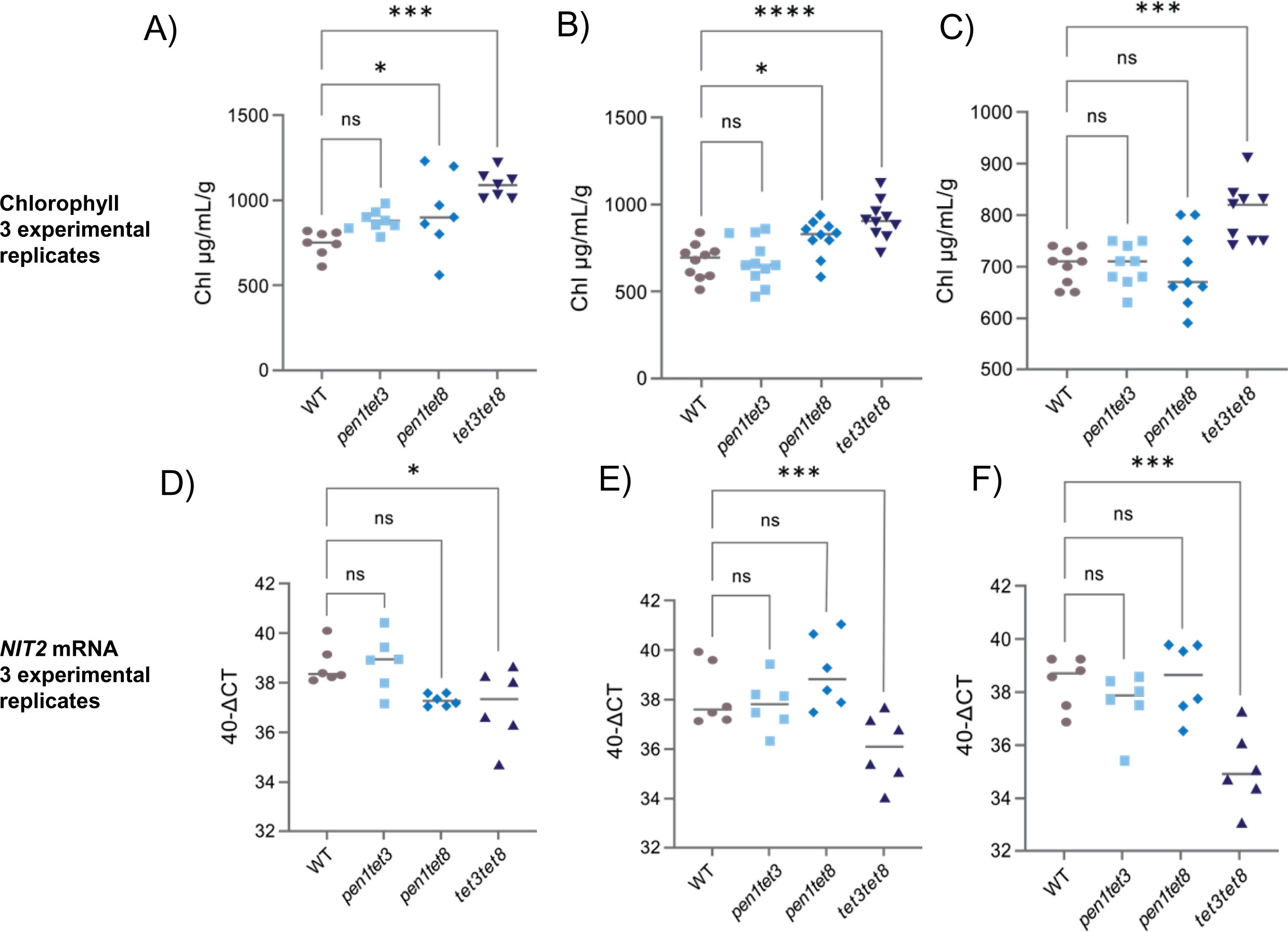
Chlorophyll and *NIT2* expression quantified after six weeks of growth in WT and double mutants. A-C) Three independent experimental replicates showing chlorophyll levels from leaf 3 (n = 6-10). D-F) *NIT2* expression in leaves 4 & 5 (n = 6). Mutations in *tet3* and *tet8* reversed the early leaf senescence observed in *pen1* while *tet3tet8* double mutants displayed a significant delay in leaf senescence.

**Figure 6.**
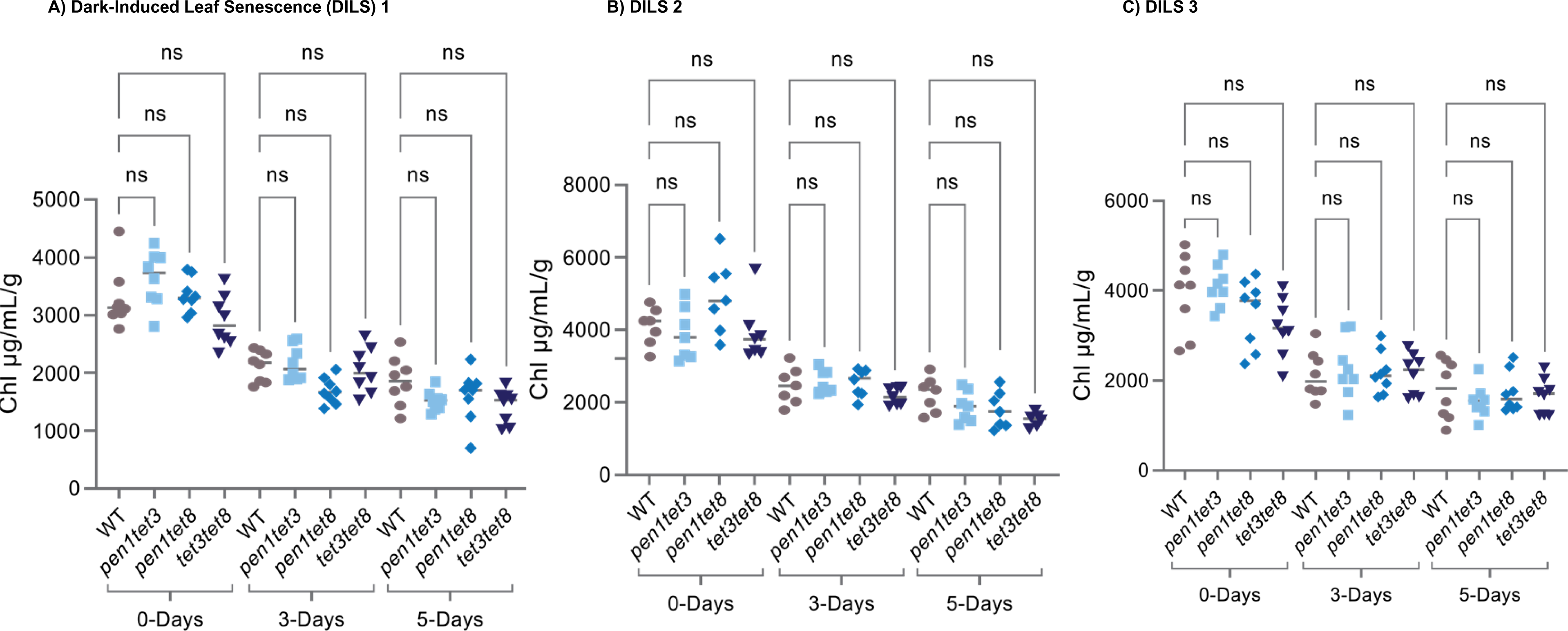
Dark induced leaf senescence in double mutants. A-C) Chlorophyll levels in three independent experiments using detached leaves treated in the dark for 0, 3 and 5 days (n = 7). There were no significant differences in chlorophyll levels at each day of dark treatment among genotypes.

## Discussion

We have presented data that positively correlate apoplast TET8 and age-related leaf senescence. Apoplast TET8 is highly abundant in early senescing *pen1* and *pen1pen3* mutants (Figure 1), and more abundant in WT as rosettes age (Figure 2). High levels of TET8 are not observed in pre-senescent *pen1* indicating that increased apoplast TET8 is not constitutively compensating for the loss of PEN1 (Figure 3), rather its high abundance coincides with leaf senescence. Further support for the correlation between apoplast TET8 and leaf senescence is the higher level in the early senescing *myb59* mutant (Figure 4). As SA has been shown to promote age-dependent leaf senescence and to increase EVs, these correlative data are not unexpected. Our mutant analysis provides genetic evidence supporting a causal relationship (Figure 5). We found that *tet8*, in combination with *tet3*, displayed a strong delayed senescence phenotype in soil-grown plants undergoing age-related senescence, but no change in DILS (Figure 6). This contrasts with *pen1* (and *pen1pen3*), which show early senescence. The genetic data support TET3 and TET8 promoting leaf senescence while PEN1 and PEN3 prevent leaf senescence. TET8 was previously found to be necessary for a full ROS and cell death response to SA (Liu et al., 2020), consistent with our results. Our model for these proposed roles is illustrated in Figure 7.

**Figure 7:**
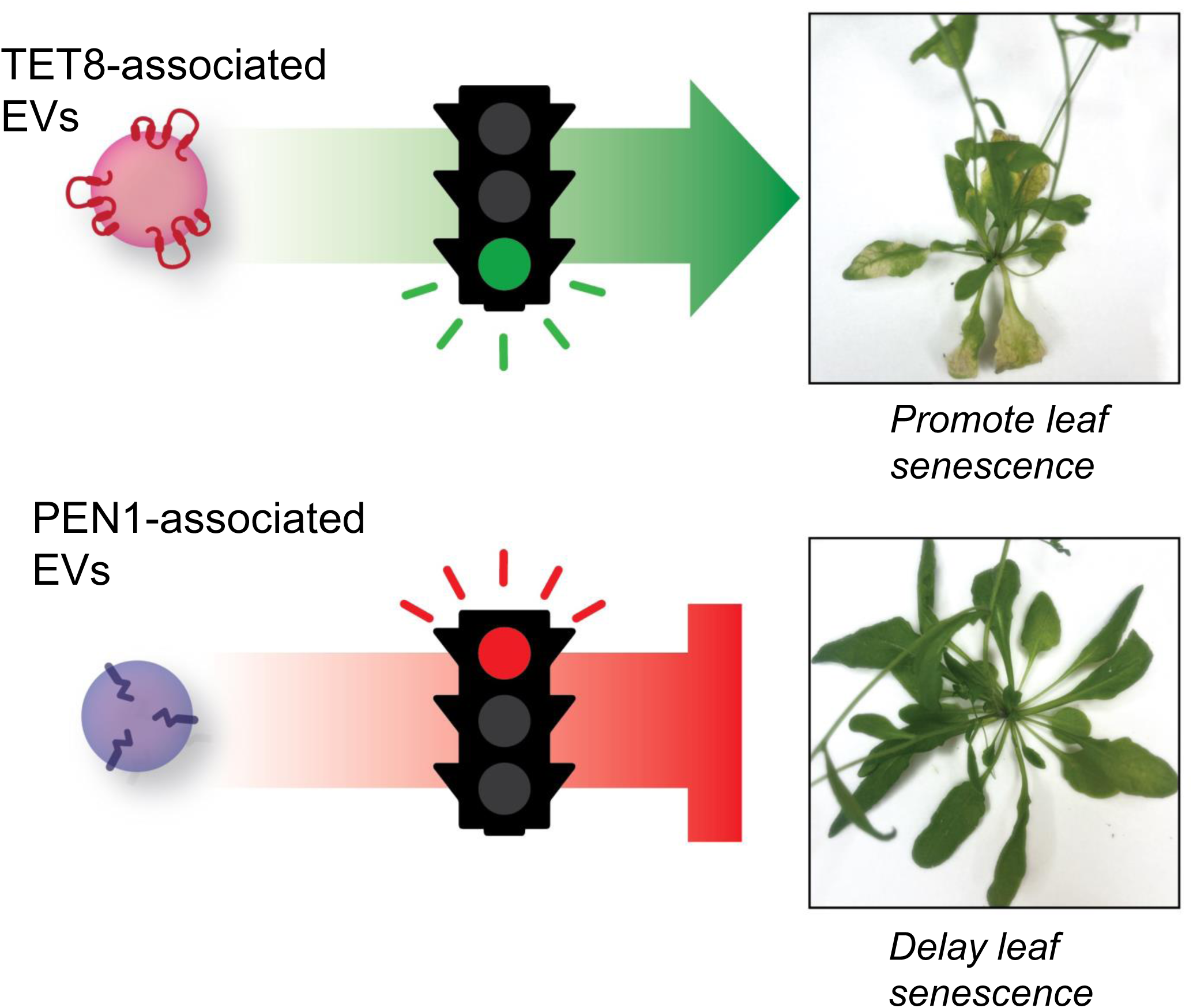
Proposed Model for EV Regulation of Leaf Senescence. TET8 and TET3 EVs localized in the apoplast promote leaf senescence in soil-grown plants and *tet3tet8* double mutants show a delay in leaf senescence. PEN1-associated EVs slow leaf senescence and *pen1* and *pen1pen3* mutants show accelerated leaf senescence.

This work measured TET8 by immunoblot in partially purified apoplast fluid that had large vesicles removed by a 10,000 x g spin. The apoplast was enriched for TET8 when compared to whole leaf extract. The TET8 signal likely arose from EVs, although plasma membrane fragments that formed into vesicles cannot be ruled out. Purifying EVs is challenging, and the current published Arabidopsis EV proteomes share unexpected proteins such as phototropin1 and phototropin2 (flavin-binding blue light photoreceptors, Christie, 2007)) that could be EV resident or from plasma membrane micelles. Current published EV proteomes also contain contaminating chloroplast proteins (Rutter and Innes, 2017; He et al., 2021), including the notoriously abundant Rubisco large subunit. This protein was most distinct in Coomassie-stained apoplast SDS-PAGE and although not the ideal candidate, was used by us and others (He et al., 2021) to compare protein amounts in apoplast samples.

When apoplast proteomes from non-senescent and senescent Arabidopsis leaves were compared, proteins related to stress-response were found to be enriched in the senescent apoplast (Borniego et al., 2020). Antifungal proteins, peroxidases, and enzymes for the catalysis of RNA were more abundant in the senescent apoplast. A few proteins that were more abundant in non-senescent leaves were shared with the EV proteome, but neither tetraspanins nor syntaxins were identified in the non-senescent or senescent apoplast proteomes. Differences in apoplast isolation (VIB versus deionized water and differences in centrifugation steps) may account for this discrepancy.

### How might tetraspanins regulate leaf senescence?

Tetraspanins are plasma membrane and endoplasmic reticulum localized proteins that form tetraspanin-enriched microdomains: dynamic hetero oligomeric protein platforms that regulate cell signaling, vesicle formation and adhesion (Reimann et al., 2017; Jimenez-Jimenez et al., 2019; Konstantinova et al., 2024; Qin et al., 2024). There are 21 tetraspanins encoded by the Arabidopsis genome (Supplemental Figure 1). A plant-specific domain has been noted in the large extracellular loop in TET1-TET13 (GCCK/RP) while TET14-17 and TOM2A-TOM2AH3 have GCC, VCC or YCC in this same position (Fujisaki et al., 2008; Boavida et al., 2013). TET18/TOM2AH2 was identified in one Arabidopsis EV proteome (Rutter and Innes, 2017). Since tetraspanins form heterodimers, the presence of multiple tetraspanins in EVs would be expected. Tetraspanins show variable expression in different plant tissues and in response to different stresses (Boavida et al., 2013; Wang et al., 2015; Qin et al., 2024)). Genetic studies reveal roles for some tetraspanins. TET1 is important for correct auxin distribution and *tet1* (*trn2*) mutants have abnormal cell divisions in the peripheral zone of the shoot apical meristem, disrupted cotyledon venation, severe effects on leaf development, and reduced numbers of vascular cell files in roots (Cnops et al., 2006; Chiu et al., 2007; Konstantinova et al., 2024). *tet5tet6* double mutants had larger leaves with greater numbers of cells suggesting redundant roles in attenuating cell division (Wang et al., 2015), while *tet3* mutants show reduced viral cell-to-cell movement via plasmodesmata (Zhu et al., 2022). Mutations in all four TOM2A-related tetraspanins show a severe growth defect (Fujisaki et al., 2008). Rice tetraspanin mutants show reduced height, reduced secondary branching of panicles and lower grain yield (Qin et al., 2024). The numerous developmental and physiological roles ascribed to tetraspanins do not provide information on mechanisms that could help define a role in regulating leaf senescence. Studies that identify TET8 and TET3 binding partners, and whether these two tetraspanins interact, may define a functional tetraspanin-enriched microdomain.

### How might PEN1 regulate leaf senescence?

PEN1 (SYP121) is a plasma membrane syntaxin or Qa-SNARE (soluble N-ethylmaleimide-sensitive factor attachment receptor). SNAREs mediate membrane fusion by forming trans-SNARE complexes ((Fujiwara et al., 2014)). Proteins secreted by PEN1 act in lipid metabolism, protein folding and cell wall modification (Waghmare et al., 2018). PEN1 localizes detergent-resistant microdomains (Qi et al., 2011) and it interacts with annexin 4, SYP71 and a hypersensitive induced reaction protein (HIR2, AT3G01290, Fujiwara et al., 2014). PEN1 is needed for the timely formation of the pre-invasive papillae that defend against haustorium-forming powdery mildew and rust fungi. H_2_O_2_-containing vesicles produced in the vicinity of pre-invasive papillae are not observed in *pen1* mutants (Collins et al., 2003) and *pen1* mutants show greater susceptibility to initial fungal penetration. A more complete pre-invasive papillae and post-invasive encasement defense is mounted in combination with SYP122, a closely related paralog. Together, these two syntaxins mediate defense against a wider range of fungi and an oomycete (Rubiato et al., 2022). The *pen1syp122* double mutant also shows a strong auto-immune response that is reversed by mutation in *FMO1*, which blocks the synthesis of N-hydroxypipecolic acid, the mobile SA signal (Zhang et al., 2007; Zhang et al., 2008). It is likely that early senescence in *pen1* and *pen1pen3* is related to autoimmunity since it was reversed with a *sid2* mutation, which reduces SA biosynthesis (Crane et al., 2019). The *pen1syp122* auto-immune response begins about two weeks after germination while the response in *pen1pen3* is weaker and commences later, after six weeks of growth (Crane et al., 2019; Rubiato et al., 2022). These observations support PEN1 negatively regulating leaf senescence. PEN1’s prevention of leaf senescence may be more related to its auto-immunity functions, potentially through its binding to HIR2, and less related to its roles in promoting pre-invasive papillae and post-invasive encasements.

Overall, we report that apoplast TET8 positively correlates to leaf senescence and that *tet3tet8* double mutants significantly delay leaf senescence. Genetic analysis suggests PEN1 slows leaf senescence, potentially related to its roles in autoimmunity. These observations support TET8-associated EVs promoting leaf senescence.

## Supplemental Figure Legends

**Supplemental Figure 1:**
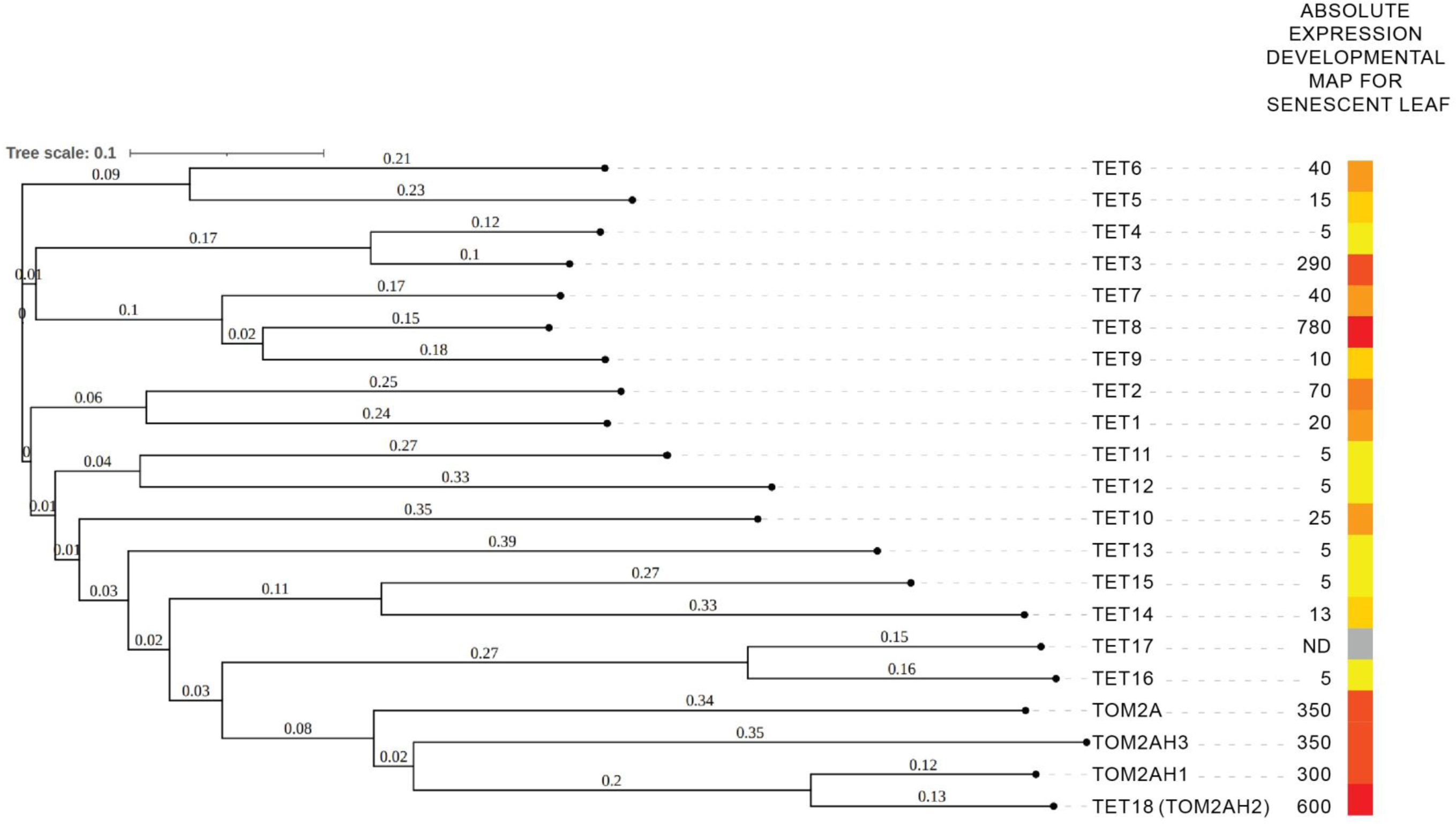
Arabidopsis Tetraspanin Phylogenetic Tree. The 21 Arabidopsis tetraspanin genes are organized in a phylogenetic tree produced using Interactive Tree of Life (iTOL). https://itol.embl.de/ The numbers adjacent to the squares on the left show the absolute expression for a senescent leaf as displayed on the Developmental Map (eFP Browser via TAIR). TET1 (AT5G46700), TET2 (AT2G19580), TET3 (AT3G45600), TET4 (AT5G60220), TET5 (AT4G23410), TET6 (AT3G12090), TET7 (AT4G28050), TET8 (AT2G23810), TET9 (AT4G30430), TET10 (AT1G63260), TET11 (AT1G18520), TET12 (AT5G23030), TET13 (AT2G03840), TET14 (AT2G01960), TET15 (AT5G57810), TET16 (AT1G18510), TET17 (AT1G74045), TET18/TOM2AH2 (AT2G20230), TOM2A (AT1G32400), TOM2AH1 (AT4G28770), TOM2AH3 (AT2G20740)

**Supplemental Figure 2:**
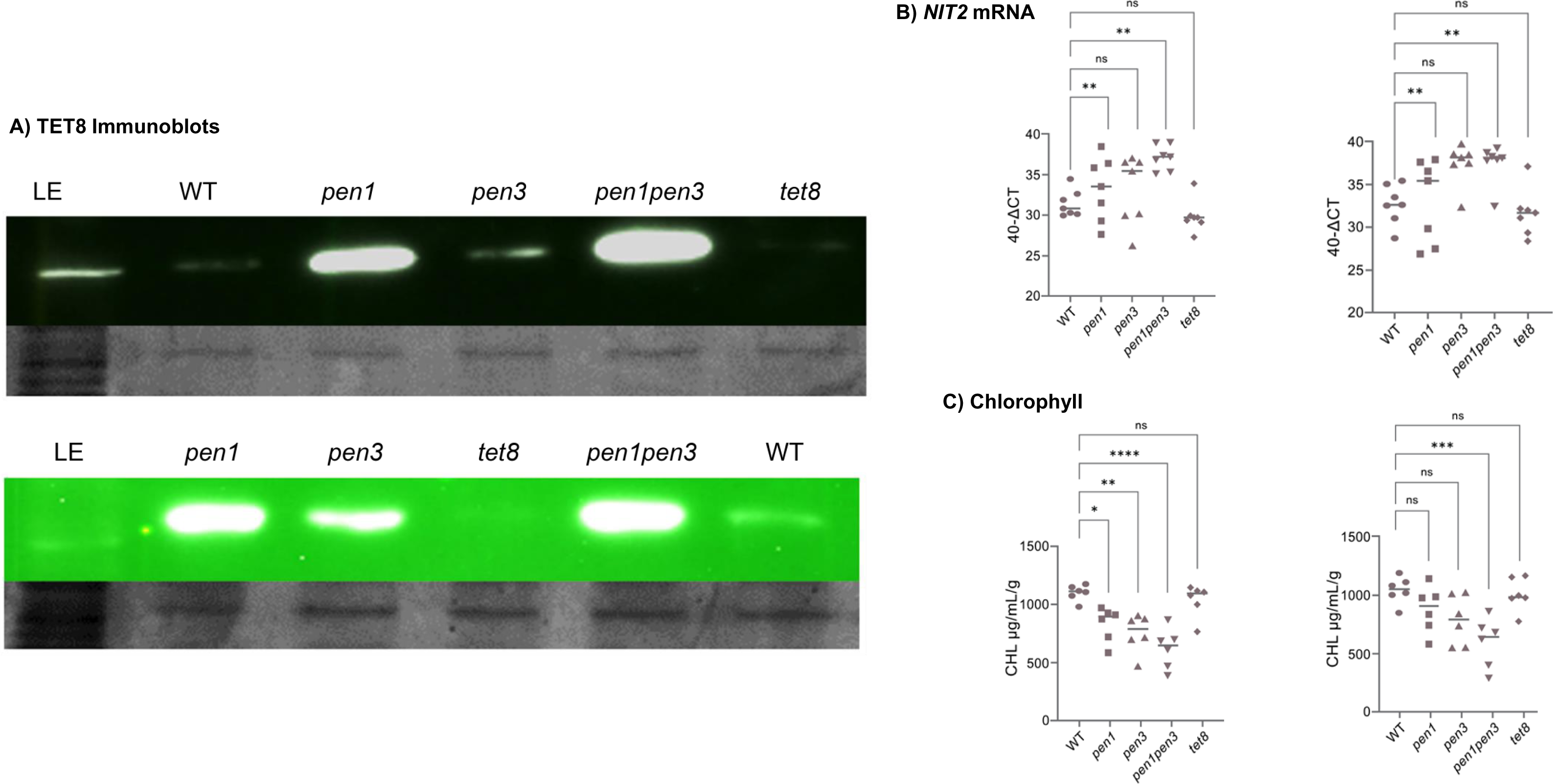
**Additional replicates showing apoplast TET8 increases in early leaf senescence mutants *pen1* and *pen1pen3***. A) TET8 was detected in leaf extract (LE) from WT and apoplast fluid from WT, *pen1*, *pen3*, *pen1pen3* and *tet8* using an immunoblot normalized to Coomassie-stained RbcL. B) From the same 8-week-old tissue, leaves 4 and 5 were harvested for RNA extraction, and *NIT2* gene expression, normalized to *ACT2*, was quantified. C) Leaf 3 was harvested, and total chlorophyll was measured and normalized to fresh weight.

**Supplemental Figure 3:**
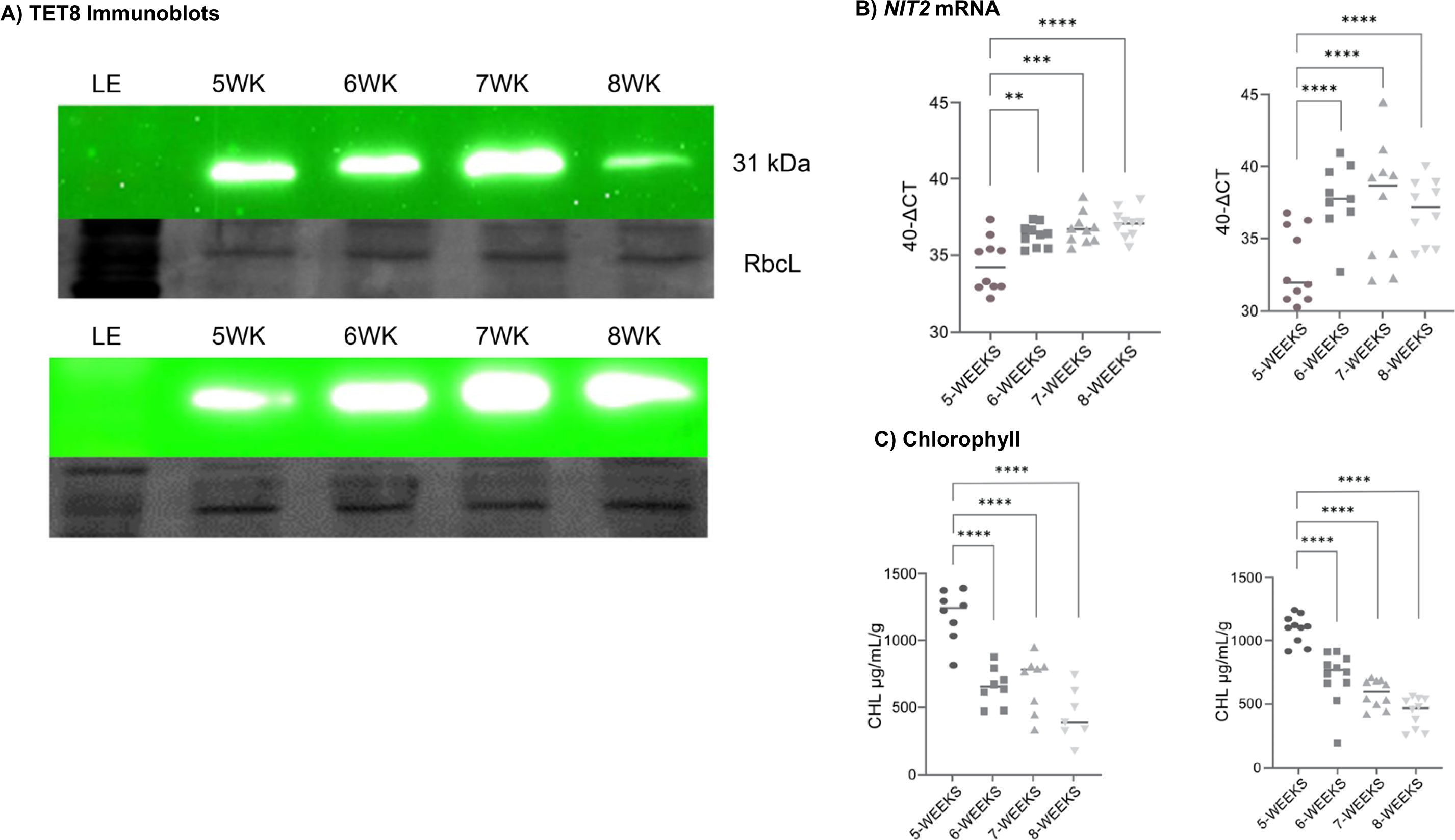
Additional replicates showing an increase in apoplast TET8 were observed in older WT rosettes. A) Apoplast TET8 was detected from rosettes harvested at 5 to 8 weeks using an immunoblot normalized to Coomassie-stained RbcL B) *NIT2* gene expression from leaves 4 and 5, and C) chlorophyll from leaf 3, are shown from plants grown alongside those used for apoplast extraction.

**Supplemental Figure 4.**
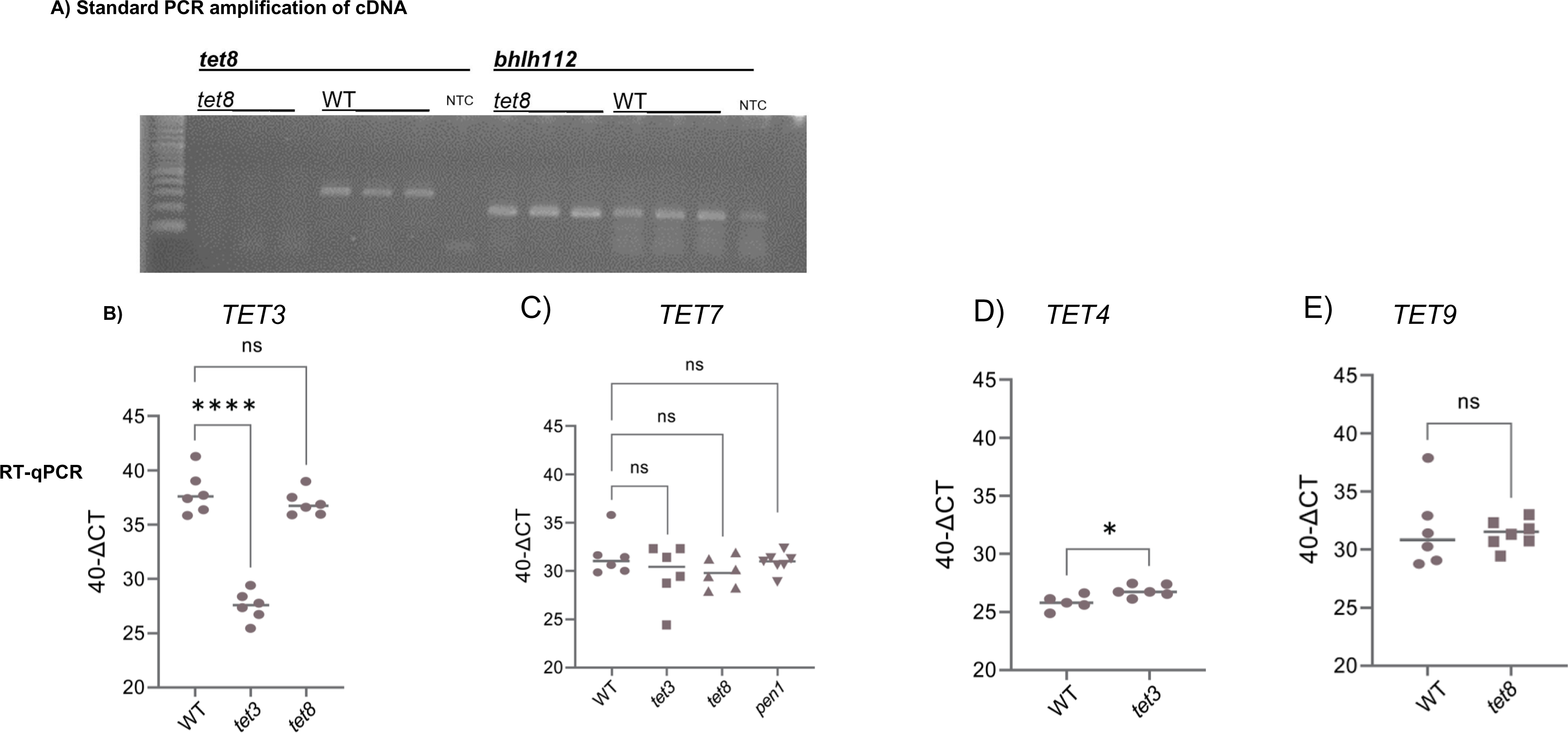
T***E***T **gene expression**. A) Full-length *TET8* mRNA cannot be amplified from *tet8* cDNA; *bHLH112* serves as a positive control. RNA was isolated from leaves from three *tet8* and three WT plants. B) *TET3* mRNA was measured by RT-qPCR and reduced by 1000-fold (2^-10) in the *tet3* mutant and unchanged in *tet8*. The 40-ΔCT value for WT levels of *TET3* and *TET8* (Figure 3C) were nearly identical. C) *TET7* mRNA levels were similar in WT, *tet3*, *tet8* and *pen1*. D) *TET4*, the most similar paralog to *TET3*, was slightly increased in *tet3*. Note that *TET4* transcript levels are ∼1000-fold lower (2^-10) than *TET3*. E) *TET9*, the most similar paralog to *TET8*, was unchanged in *tet8*. *TET9* expression is ∼64-fold lower (2^-6) than *TET8*. For panels B-E, RNA was isolated from leaves 4 & 5 after 8 weeks of growth (n = 6). One outlier was removed from WT *TET4* by outlier tests in GraphPad Prism, no other outliers were identified.

## Acknowledgements

This work was supported by NIH RISE grant R25GM071638 and Small Faculty Grants from California State University, Long Beach. We would like to thank Will Hinckley and Michelle Smith for critical reading of the manuscript.

## Author Contributions

JZ, BV and AHH designed and performed the research, developed and improved protocols, and contributed to the quantitative analysis of their data. JB designed the research, contributed to the statistical analysis, and wrote the paper. All authors were involved in the editing process.

---

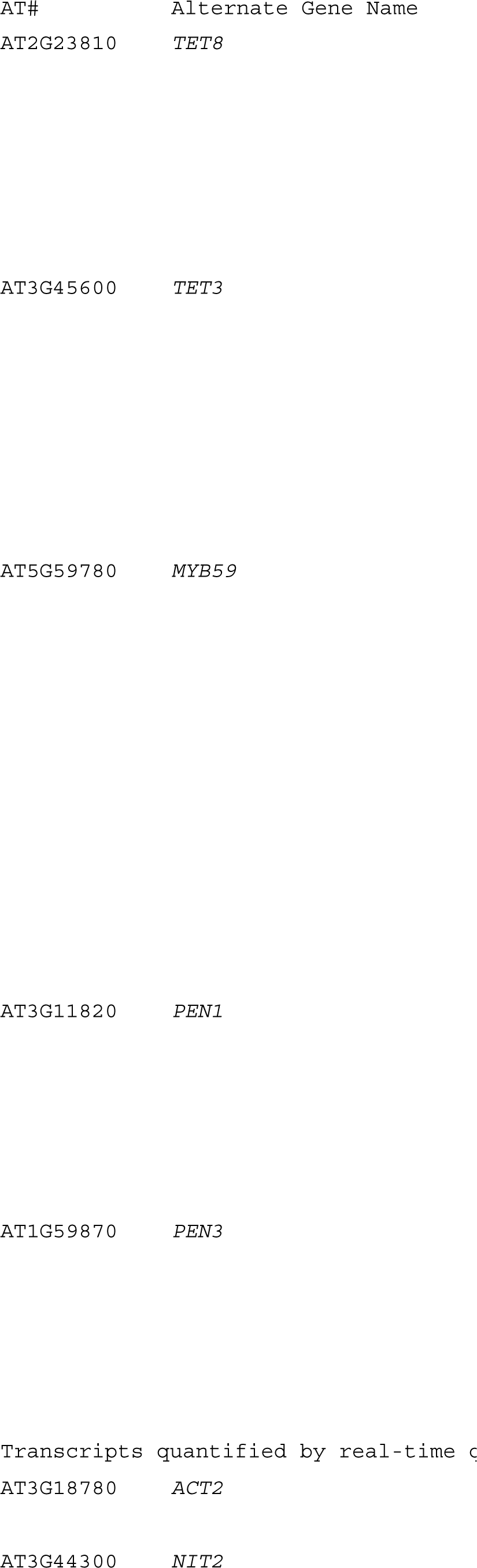
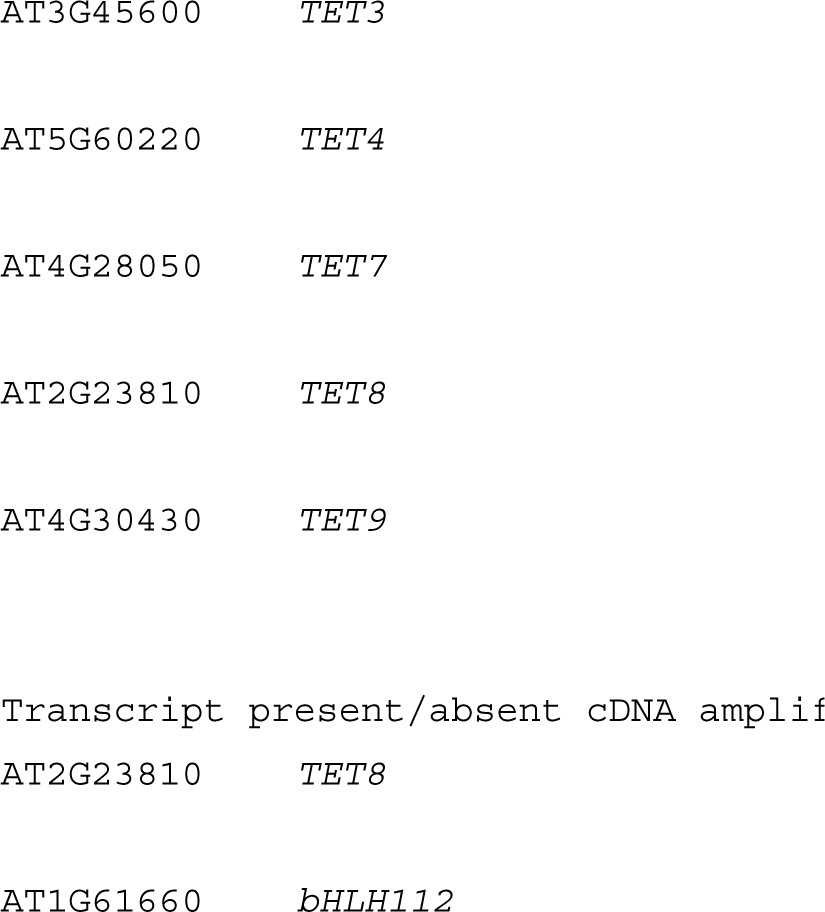
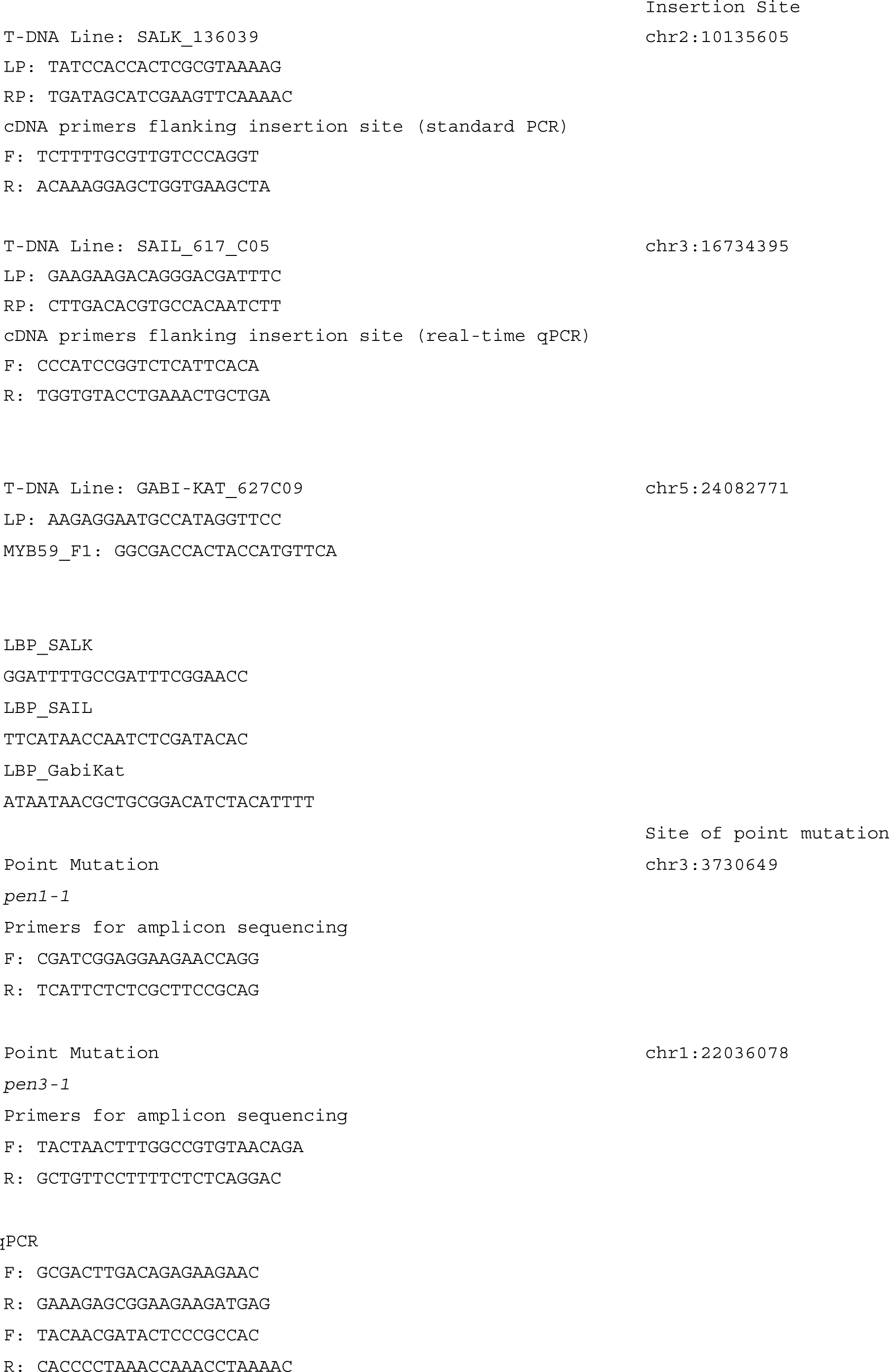
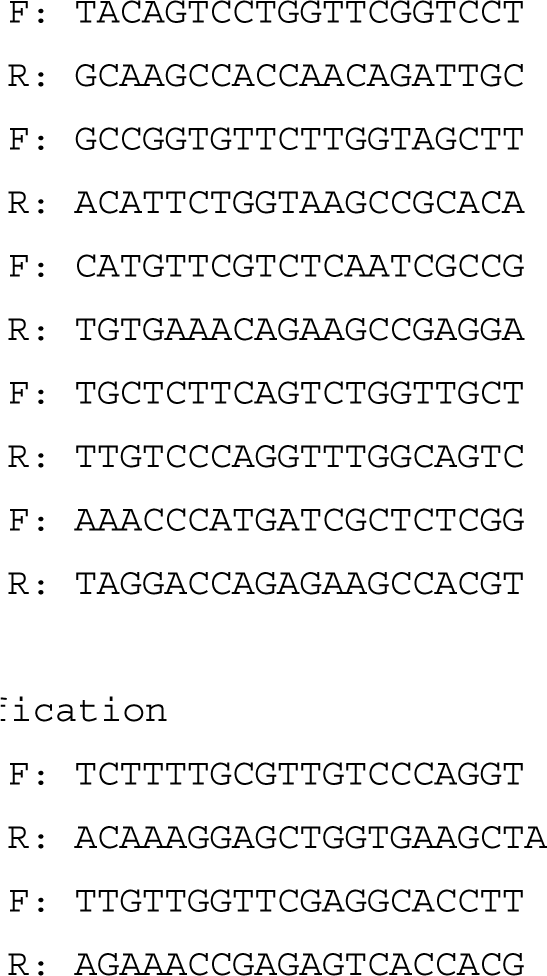
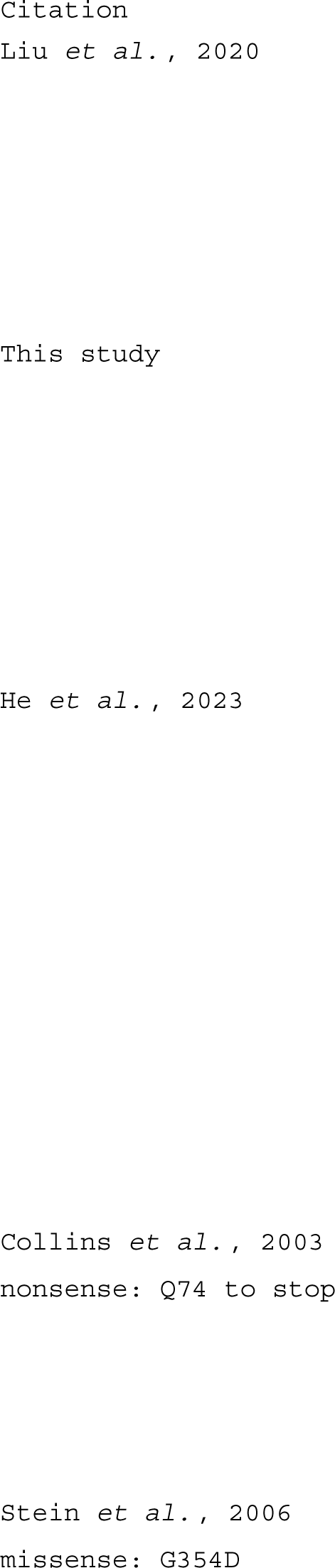

## References

An Q, Ehlers K, Kogel KH, Van Bel AJE, Hückelhoven R (2006a) Multivesicular compartments proliferate in susceptible and resistant MLA12-barley leaves in response to infection by the biotrophic powdery mildew fungus. New Phytologist 172: 563–576

An Q, Huckelhoven R, Kogel K-H, van Bel AJE (2006b) Multivesicular bodies participate in a cell wall-associated defence response in barley leaves attacked by the pathogenic powdery mildew fungus. Cell Microbiol 8: 1009–1019

Baldrich P, Rutter BD, Karimi HZ, Podicheti R, Meyers BC, Innes RW (2019) Plant Extracellular Vesicles Contain Diverse Small RNA Species and Are Enriched in 10- to 17-Nucleotide “Tiny” RNAs. Plant Cell. doi: 10.1105/tpc.18.00872

Boavida LC, Qin P, Broz M, Becker JD, McCormick S (2013) Arabidopsis Tetraspanins Are Confined to Discrete Expression Domains and Cell Types in Reproductive Tissues and Form Homo- and Heterodimers When Expressed in Yeast . Plant Physiol 163: 696–712

Borniego ML, Molina MC, Guiamét JJ, Martinez DE (2020) Physiological and Proteomic Changes in the Apoplast Accompany Leaf Senescence in Arabidopsis. Front Plant Sci. doi: 10.3389/fpls.2019.01635

Bose S, Aggarwal S, Singh DV, Acharya N (2020) Extracellular vesicles: An emerging platform in gram-positive bacteria. Microbial Cell 7: 312–322

Brusslan JA, Bonora G, Rus-Canterbury AM, Tariq F, Jaroszewicz A, Pellegrini M (2015) A Genome-Wide Chronological Study of Gene Expression and Two Histone Modifications, H3K4me3 and H3K9ac, during Developmental Leaf Senescence. Plant Physiol 168: 1246– 1261

Brusslan JA, Rus Alvarez-Canterbury AM, Nair NU, Rice JC, Hitchler MJ, Pellegrini M (2012) Genome-wide evaluation of histone methylation changes associated with leaf senescence in Arabidopsis. PLoS One 7: e33151

Buchanan-Wollaston V, Page T, Harrison E, Breeze E, Lim PO, Nam HG, Lin J-F, Wu S-H, Swidzinski J, Ishizaki K, et al (2005) Comparative transcriptome analysis reveals significant differences in gene expression and signalling pathways between developmental and dark/starvation-induced senescence in Arabidopsis. The Plant Journal. doi: 10.1111/j.1365-313X.2005.02399.x

Cai Q, Qiao L, Wang M, He B, Lin F-M, Palmquist J, Huang S-D, Jin H (2018) Plants send small RNAs in extracellular vesicles to fungal pathogen to silence virulence genes. Science (1979) 360: 1126–1129

Chaya T, Banerjee A, Rutter BD, Adekanye D, Ross J, Hu G, Innes RW, Caplan JL (2024) The extracellular vesicle proteomes of *Sorghum bicolor* and *Arabidopsis thaliana* are partially conserved. Plant Physiol 194: 1481–1497

Chiu W-H, Chandler J, Cnops G, Van Lijsebettens M, Werr W (2007) Mutations in the TORNADO2 gene affect cellular decisions in the peripheral zone of the shoot apical meristem of Arabidopsis thaliana. Plant Mol Biol 63: 731–744

Christie JM (2007) Phototropin Blue-Light Receptors. Annu Rev Plant Biol 58: 21–45

Cnops G, Neyt P, Raes J, Petrarulo M, Nelissen H, Malenica N, Luschnig C, Tietz O, Ditengou F, Palme K, et al (2006) The *TORNADO1* and *TORNADO2* Genes Function in Several Patterning Processes during Early Leaf Development in *Arabidopsis thaliana*. Plant Cell 18: 852–866

Collins NC, Thordal-Christensen H, Lipka V, Bau S, Kombrink E, Qiu J-L, Huckelhoven R, Stein M, Freialdenhoven A, Somerville S, et al (2003) SNARE-protein-meidated disease resistance at the plant cell wall. Nature 425: 973–977

Crane RA, Cardénas Valdez M, Castaneda N, Jackson CL, Riley CJ, Mostafa I, Kong W, Chhajed S, Chen S, Brusslan JA (2019) Negative Regulation of Age-Related Developmental Leaf Senescence by the IAOx Pathway, PEN1, and PEN3. Front Plant Sci. doi: 10.3389/fpls.2019.01202

Dai J, Su Y, Zhong S, Cong L, Liu B, Yang J, Tao Y, He Z, Chen C, Jiang Y (2020) Exosomes: key players in cancer and potential therapeutic strategy. Signal Transduct Target Ther 5: 145

Fujisaki K, Kobayashi S, Tsujimoto Y, Naito S, Ishikawa M (2008) Analysis of tobamovirus multiplication in Arabidopsis thaliana mutants defective in TOM2A homologues. Journal of General Virology 89: 1519–1524

Fujiwara M, Uemura T, Ebine K, Nishimori Y, Ueda T, Nakano A, Sato MH, Fukao Y (2014) Interactomics of Qa-SNARE in Arabidopsis thaliana. Plant Cell Physiol 55: 781–789

Garapati P, Xue G-P, Munné-Bosch S, Balazadeh S (2015) Transcription Factor ATAF1 in Arabidopsis Promotes Senescence by Direct Regulation of Key Chloroplast Maintenance and Senescence Transcriptional Cascades. Plant Physiol 168: 1122–1139

He B, Cai Q, Qiao L, Huang C-Y, Wang S, Miao W, Ha T, Wang Y, Jin H (2021) RNA-binding proteins contribute to small RNA loading in plant extracellular vesicles. Nat Plants 7: 342– 352

He S, Zhi F, Min Y, Ma R, Ge A, Wang S, Wang J, Liu Z, Guo Y, Chen M (2023) The MYB59 transcription factor negatively regulates salicylic acid- and jasmonic acid-mediated leaf senescence. Plant Physiol 192: 488–503

He Y, Xu J, Wang X, He X, Wang Y, Zhou J, Zhang S, Meng X (2019) The Arabidopsis Pleiotropic Drug Resistance Transporters PEN3 and PDR12 Mediate Camalexin Secretion for Resistance to *Botrytis cinerea*. Plant Cell 31: 2206–2222

Heard W, Sklenář J, Tomé DFA, Robatzek S, Jones AME (2015) Identification of Regulatory and Cargo Proteins of Endosomal and Secretory Pathways in Arabidopsis thaliana by Proteomic Dissection*. Molecular & Cellular Proteomics 14: 1796–1813

Jeppesen DK, Fenix AM, Franklin JL, Higginbotham JN, Zhang Q, Zimmerman LJ, Liebler DC, Ping J, Liu Q, Evans R, et al (2019) Reassessment of Exosome Composition. Cell 177: 428–445.e18

Jimenez-Jimenez S, Hashimoto K, Santana O, Aguirre J, Kuchitsu K, Cárdenas L (2019) Emerging roles of tetraspanins in plant inter-cellular and inter-kingdom communication. Plant Signal Behav 14: e1581559

Konstantinova N, Mor E, Verhelst E, Nolf J, Vereecken K, Wang F, Van Damme D, De Rybel B, Glanc M (2024) A precise balance of <SCP>TETRASPANIN1</SCP> / <SCP>TORNADO2</SCP> activity is required for vascular proliferation and ground tissue patterning in Arabidopsis. Physiol Plant. doi: 10.1111/ppl.14182

Lananna B V., Imai S (2021) Friends and foes: Extracellular vesicles in aging and rejuvenation. FASEB Bioadv 3: 787–801

Liebana-Jordan M, Brotons B, Falcon-Perez JM, Gonzalez E (2021) Extracellular Vesicles in the Fungi Kingdom. Int J Mol Sci 22: 7221

Liu N-J, Wang N, Bao J-J, Zhu H-X, Wang L-J, Chen X-Y (2020) Lipidomic Analysis Reveals the Importance of GIPCs in Arabidopsis Leaf Extracellular Vesicles. Mol Plant 13: 1523– 1532

Livak KJ, Schmittgen TD (2001) Analysis of relative gene expression data using real-time quantitative PCR and the 2(-Delta Delta C(T)) Method. Methods 25: 402–408

Lu X, Dittgen J, Pislewska-Bednarek M, Molina A, Schneider B, Svatos A, Doubsky J, Schneeberger K, Weigel D, Bednarek P, et al (2015) Mutant allele-specific uncoupling of penetration3 functions reveals engagement of the ATP-binding cassette transporter in distinct tryptophan metabolic pathways. Plant Physiol. doi: 10.1104/pp.15.00182

Porra RJ, Thompson WA, Kriedemann PE (1989) Determination of accurate extinction coefficients and simultaneous equations for assaying chlorophylls a and b extracted with four different solvents: verification of the concentration of chlorophyll standards by atomic absorption spectroscopy. Biochim Biophys Acta 975: 384–394

Qi Y, Tsuda K, Nguyen L V., Wang X, Lin J, Murphy AS, Glazebrook J, Thordal-Christensen H, Katagiri F (2011) Physical Association of Arabidopsis Hypersensitive Induced Reaction Proteins (HIRs) with the Immune Receptor RPS2. Journal of Biological Chemistry 286: 31297–31307

Qin S, Li W, Zeng J, Huang Y, Cai Q (2024) Rice tetraspanins express in specific domains of diverse tissues and regulate plant architecture and root growth. The Plant Journal 117: 892–908

Regente M, Corti-Monzón G, Maldonado AM, Pinedo M, Jorrín J, de la Canal L (2009) Vesicular fractions of sunflower apoplastic fluids are associated with potential exosome marker proteins. FEBS Lett 583: 3363–3366

Reimann R, Kost B, Dettmer J (2017) TETRASPANINs in Plants. Front Plant Sci. doi: 10.3389/fpls.2017.00545

Rubiato HM, Liu M, O’Connell RJ, Nielsen ME (2022) Plant SYP12 syntaxins mediate an evolutionarily conserved general immunity to filamentous pathogens. Elife. doi: 10.7554/eLife.73487

Rutter B, Rutter K, Innes R (2017) Isolation and Quantification of Plant Extracellular Vesicles. Bio Protoc. doi: 10.21769/BioProtoc.2533

Rutter BD, Innes RW (2017) Extracellular Vesicles Isolated from the Leaf Apoplast Carry Stress-Response Proteins. Plant Physiol 173: 728–741

Stein M, Diottgen J, Sanchez-Rodriguez C, Hou B-H, Molina A, Schulze-Lefert P, Lipka V, Somerville S (2006) Arabidopsis PEN3/PDR8, an ATP Binding Cassette Transporter, Contributes to Nonhost Resistance to Inappropriate Pathogens That Enter by Direct Penetration. THE PLANT CELL ONLINE 18: 731–746

Waghmare S, Lileikyte E, Karnik R, Goodman JK, Blatt MR, Jones AME (2018) SNAREs SYP121 and SYP122 Mediate the Secretion of Distinct Cargo Subsets. Plant Physiol 178: 1679–1688

Wang F, Muto A, Van de Velde J, Neyt P, Himanen K, Vandepoele K, Van Lijsebettens M (2015) Functional Analysis of Arabidopsis TETRASPANIN Gene Family in Plant Growth and Development. Plant Physiol pp.01310.2015

Wang S, He B, Wu H, Cai Q, Ramírez-Sánchez O, Abreu-Goodger C, Birch PRJ, Jin H (2024) Plant mRNAs move into a fungal pathogen via extracellular vesicles to reduce infection. Cell Host Microbe 32: 93–105.e6

Zand Karimi H, Baldrich P, Rutter BD, Borniego L, Zajt KK, Meyers BC, Innes RW (2022) Arabidopsis apoplastic fluid contains sRNA- and circular RNA–protein complexes that are located outside extracellular vesicles. Plant Cell 34: 1863–1881

Zhang Z, Feechan A, Pedersen C, Newman M, Qiu J, Olesen KL, Thordal-Christensen H (2007) A SNARE-protein has opposing functions in penetration resistance and defence signalling pathways. The Plant Journal 49: 302–312

Zhang Z, Lenk A, Andersson MX, Gjetting T, Pedersen C, Nielsen ME, Newman M-A, Hou B-H, Somerville SC, Thordal-Christensen H (2008) A Lesion-Mimic Syntaxin Double Mutant in Arabidopsis Reveals Novel Complexity of Pathogen Defense Signaling. Mol Plant 1: 510–527

Zhu T, Sun Y, Chen X (2022) Arabidopsis Tetraspanins Facilitate Virus Infection via Membrane-Recognition GCCK/RP Motif and Cysteine Residues. Front Plant Sci. doi: 10.3389/fpls.2022.805633

